# Neural computations in the foveal and peripheral visual fields during active search

**DOI:** 10.1101/2021.11.22.469359

**Authors:** Jie Zhang, Xiaocang Zhu, Zhengyu Ma, Shanshan Wang, Yutian Wang, Hossein Esteky, Yonghong Tian, Huihui Zhou

## Abstract

Active vision requires coordinated attentional processing across both foveal and peripheral receptive fields (RFs), yet the underlying neural dynamics and computational mechanisms remain poorly understood. Previous research has predominantly focused on attention in the visual periphery, leaving the role of foveal processing in naturalistic tasks largely unexplored. Here, we recorded neural activity from both foveal and peripheral RFs in areas V4 and IT of monkeys during free-gaze visual search among complex stimuli. We found robust feature-based attentional enhancements in foveal units, challenging the prevailing view that such modulation is predominantly peripheral. By integrating data from foveal and peripheral recordings, we revealed a non-uniform, dynamically distributed pattern of feature attention across the visual field. Behaviorally, foveal attentional enhancements promoted sustained or repeated fixations on targets, while peripheral attentional signals facilitated target detection and guidance of future saccades. These findings suggest that foveal and peripheral attention operate in a complementary fashion, and highlight the critical role of foveal feature attention in shaping global attention allocation and fixation behavior during active vision. This work advances our understanding of the neural computations that support complex visual search and underscores the need to account for foveal processing in models of attention.

## Introduction

Visual attention is a cornerstone of cognitive processing, enabling the brain to efficiently prioritize task-relevant visual information. Visual search has been extensively studied to understand the neural mechanisms of visual attention during active vision, which is essential for efficient information gathering (1, 2). Feature-based attention enhances the visual responses of neurons to stimuli that share features with the target in areas such as V4, IT, the lateral intraparietal cortex (LIP), and the prefrontal cortex (PFC) (3–9), and can shift the tuning of V4 neurons to more closely match the spectral properties of the target (10). These feature-based attentional effects are thought to occur in parallel across multiple locations in the visual field—a view commonly referred to as distributed feature attention (1, 3, 11–15). In contrast, overt spatial attention enhances responses to stimuli that serve as saccade targets during visual search. Spatial and feature-based attentional processes exhibit temporal overlap in V4, PFC, and LIP (5, 6, 8). Together, parallel feature-based modulation and the serial selection of saccade targets guide eye movements for efficient visual search (2, 3, 16). Notably, most of these findings have been derived from analyses of visual and attentional processing in the peripheral visual field.

A number of behavioral studies demonstrate that processing in the foveal field plays a critical role in visual search. Masking the foveal visual field severely disrupts visual exploration behavior, leading to decreased search accuracy, increased search time, and the elimination of search facilitation in repeated displays (17–20). These behavioral impairments are comparable to—or even greater than—those caused by masking the peripheral field (18, 19). Moreover, high attentional load in the foveal region reduces detection accuracy for peripheral stimuli (21), and directing attention to peripheral regions diminishes EEG responses to foveal stimuli (22). Similarly, attending to a specific feature during a central task impairs performance in the periphery (23, 24). Other studies suggest that foveal and peripheral processes can operate independently (23–26). V4 and inferotemporal cortex (IT), as the middle and high-level areas of the ventral visual stream, are important for object recognition and categorization (27–34), and their roles have been extensively studied in central vision. At the neuronal level, however, most investigations have largely neglected their functions during active, free-gaze visual search. The prefrontal cortex, including LPFC, has long been implicated as a source of top-down signals that bias the selection of attended features and modulate visual cortical responses (6, 9, 11, 35–40). Although target-related visual responses have been reported in IT during visual exploration (41), and target-selective responses have been observed in the human medial temporal lobe (MTL) (42) and medial frontal cortex (MFC) (43) during visual search, these studies did not map the receptive fields (RFs) of recorded neurons. As a result, the neural computations and dynamics underlying foveal visual attention—and how they interact with peripheral attentional processes—remain largely unknown.

Efficient visual search requires seamless coordination between foveal analysis, which supports detailed inspection of the currently fixated object, and peripheral selection, which guides saccades toward potential targets. However, most existing models of visual search focus almost exclusively on peripheral guidance, overlooking how foveal processing interacts with attentional mechanisms across the visual field. To address this gap, we sought to uncover the neuronal mechanisms that coordinate foveal and peripheral attention during goal-directed visual search. We hypothesized that feature-based attention operates concurrently in both foveal and peripheral representations but with distinct and complementary functions: foveal attention enhances the processing of the currently fixated target to sustain or reinitiate fixation, whereas peripheral attention facilitates the detection of potential targets and guides saccades toward them. To test this hypothesis, we will simultaneously record neuronal activity from units with foveal and peripheral receptive fields in areas V4, IT, and lateral prefrontal cortex (LPFC) while monkeys perform a naturalistic category-based visual search task. We will analyze attentional modulation of neural responses across visual-field locations, temporal dynamics of foveal and peripheral signals during fixation and saccade planning, and cross-area interactions linking visual and prefrontal circuits. These analyses will reveal how foveal and peripheral attention signals are distributed, coordinated, and functionally coupled to behavior. By integrating neuronal, behavioral, and population-level analyses, this study aims to refine models of attentional control by providing a unified account of how the brain flexibly balances fixation maintenance and target selection during naturalistic visual search.

## Results

Two monkeys performed a category-based visual search task, where their objective was to fixate on one of the two search targets that matched the category of the cue (**Fig. 1A, B**). Specifically, the monkeys were presented with a central fixation point for 400 ms, followed by a cue lasting 500-1300 ms. After a 500 ms delay, a search array appeared with 11 items, including two targets, randomly chosen from 20 possible locations (**Fig. 1E**). The monkeys had 4000 ms to find one target and maintain fixation on it for 800 ms to earn a juice reward. Fixating on either target completed the trial, and the monkeys did not search for the second target. A new trial began after the reward. It is worth noting that the two target stimuli matched the category of the cue but were different images. The monkeys were required to maintain fixation throughout the cue and delay periods. During search, however, eye movements were unconstrained, and monkeys could revisit each search distractor or target as long as they did not fixate on a target for 800 ms. Both monkeys performed the task proficiently: Monkey S achieved 92% accuracy, and Monkey E achieved 86%. The mean number of saccades monkeys made to find the target after the onset of the search array was 2.25 ± 1.35 (mean ± SD across trials; **Table 1**) of correct trials, and the mean saccade amplitude was 7.99° ± 3.58° (mean ± SD across saccades; **Table 1**). Monkeys could fixate on each distractor or the target freely, provided they did not maintain fixation on the target for longer than 800 ms. Across sessions, 42.44% ± 3.6% of saccades were directed to distractors, 57.56% ± 3.6% to targets, and 12.59% ± 3.46% were saccades away from targets (see our previous studies (44–46) for detailed behavioral analyses). We further examined the probability of fixation transitions as a function of fixation duration (**Fig. 1C**). Interestingly, we found that the probability of saccading to the other target decreased with increasing duration of the current target fixation. In contrast, the probability of re-fixating on the target increased with fixation duration of that target. However, the probability of saccading to a target did not vary with current distractor fixation duration.

**Fig. 1.**
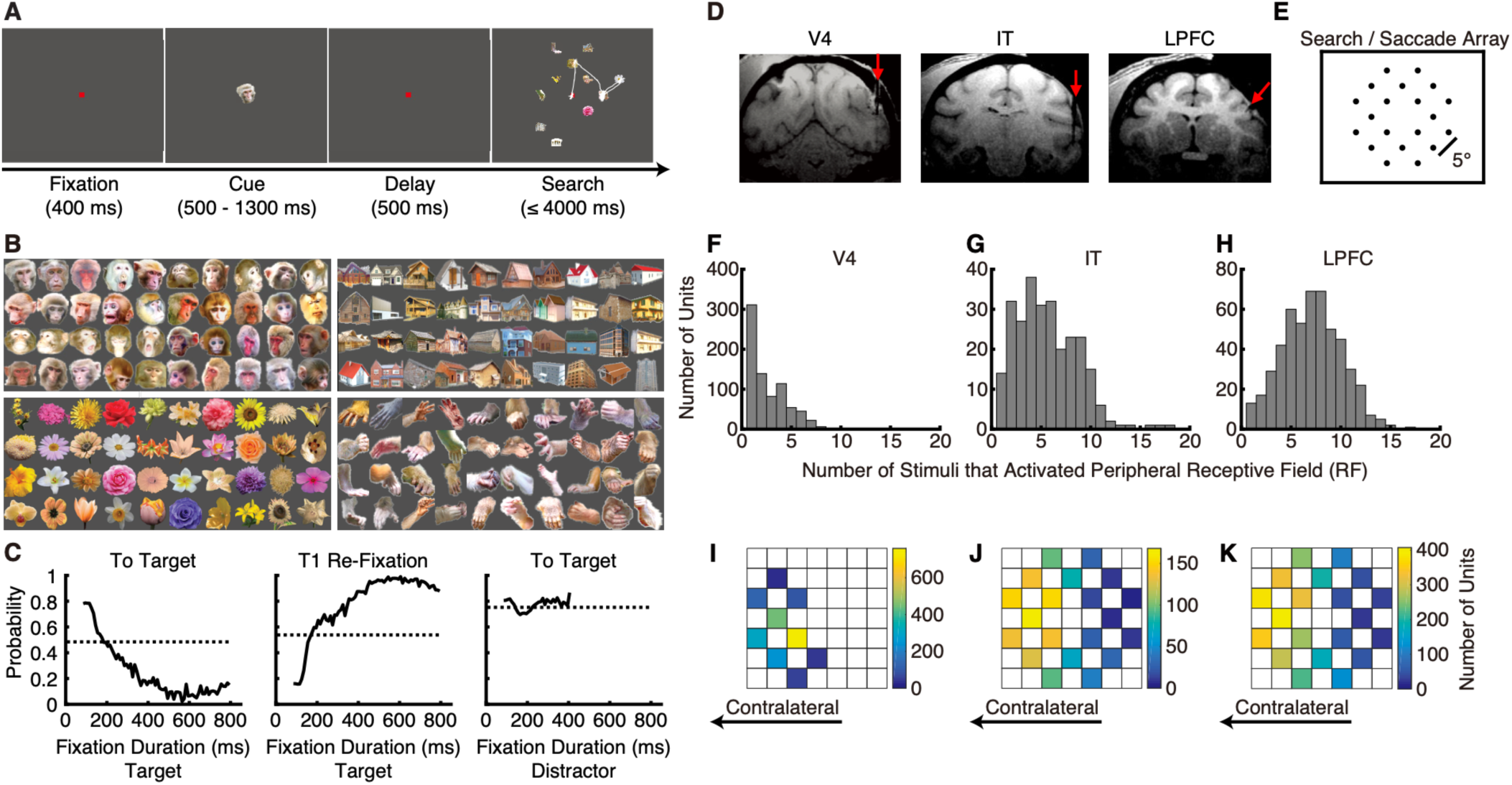
Task and recording sites. **(A)** Behavioral task. A central cue was first presented to indicate the category of the target to be searched for. This was followed by a search array consisting of eleven stimuli, including two target stimuli and nine distractors. The cue and the two targets belonged to the same category, although the targets were visually distinct from the cue. Monkeys were rewarded for maintaining fixation on either target for at least 800 ms. The white trace indicates eye gazes. In this example, the monkey started from the center, fixated on one face target for less than 800 ms, moved to another face target and fixated for less than 800 ms, then moved to a distractor, and finally shifted back to the face target, fixating for 800 ms to complete the trial. **(B)** Stimuli. Four categories of visual objects (40 images per category) were used for neural recordings. **(C)** Probability of fixation transitions as a function of fixation duration. **(D)** MRI images showing representative recording sites in areas V4, IT, and LPFC. Red arrows indicate the directions of electrode penetrations. **(E)** The 20 possible stimulus locations used in the search array. Peripheral receptive fields (RFs) were mapped using the visually guided saccade task, which had the same 20 possible stimulus locations as the visual search task. **(F-H)** Histogram of the number of stimuli that activated the peripheral RF. **(I-K)** The aggregated tuning regions of the peripheral units. Color bars show the number of units with tuning regions in a given stimulus location. The right side of the brain was recorded for both monkeys. **(F, I)** V4 units. **(G, J)** IT units. **(H, K)** LPFC units.

**Table 1.** Summary of saccade probabilities, saccade counts, and saccade amplitudes (mean ± SD).

|  | All Trial | Face Trial | House Trial |
| --- | --- | --- | --- |
| Prob(to target) | 57.56% ± 3.6% | 58.91% ± 5.24% | 56.45% ± 2.77% |
| Prob(from target) | 12.59% ± 3.46% | 12.17% ± 3.23% | 12.98% ± 3.88% |
| Prob(to/from distractor) | 42.44% ± 3.6% | 41.09% ± 5.24% | 43.55% ± 2.77% |
| Saccade Count Per Trial | 2.25 ± 1.35 | 2.19 ± 1.33 | 2.32 ± 1.36 |
| Saccade Amplitude | 7.99° ± 3.58° | 8.07° ± 3.61° | 7.91° ± 3.55° |

Neuronal activity was simultaneously recorded in areas V4, IT, and LPFC (see **Fig. 1D** for representative recording sites; see (44–47) for more details). In V4, 1898 units with foveal RFs and 765 units with peripheral RFs were recorded. In IT, 1511 foveal units and 239 peripheral units were recorded. In LPFC, 35 foveal units and 507 peripheral units were recorded. The foveal units showed significantly increased responses to the cue stimulus presented at the center of the screen (Wilcoxon rank-sum test, *P* < 0.05), but not to the appearance of the search array in the peripheral field (**Fig. S1A–C, E–G**). Notably, the items in the search array were presented at least 5° from the central fixation point and were also separated by at least 5° from each other, excluding the possibility that peripheral stimuli fell within their foveal RFs during fixations. In contrast, peripheral units responded to stimuli in the peripheral field but not to the centrally presented cue stimulus (**Fig. S1I–K, M–O, Q–S**). The RFs of these peripheral units were further mapped using a visually guided saccade task and quantified by the number of stimuli that activated each unit (**Fig. 1F-K**). The eccentricities of the peripheral RFs were 6.22° ± 1.31° (mean ± SD) in V4, 7.04° ± 1.52° in IT, and 6.68° ± 1.56° in LPFC. The sizes of the peripheral RFs were 3.67° ± 1.87° in V4, 6.86° ± 3.11° in IT, and 8.65° ± 3.02° in LPFC. The numbers of items from the search array falling within peripheral RFs were 1.49 ± 0.55 in V4, 2.2 ± 0.72 in IT, and 2.56 ± 0.74 in LPFC (also see our previous study (44)). All further analyses were based on these foveal and peripheral units. Results were qualitatively similar in both monkeys and were therefore combined.

### Feature attentional modulation in V4 and IT foveal units during visual search

We first assessed the category selectivity of units using a selectivity index (see **Methods**), following methodologies similar to those used in previous studies (48, 49). Units were categorized as face-selective or house-selective if their mean responses to the two categories were statistically different and the mean response to the preferred category was at least 130% of the response to the non-preferred category, with selectivity index values greater than 0.13 or less than −0.13 (see **Methods**). Units that did not exhibit a significant difference in their mean responses to face versus house stimuli were classified as non-selective. Among the 1898 V4 foveal units, 266 were categorized as face-selective, 304 as house-selective, and 1051 as non-selective. Of the 1511 IT foveal units, 518 were face-selective, 340 were house-selective, and 558 were non-selective (see **Fig. S1D, H** for the distributions of selectivity index for all foveal units in V4 and IT).

All analyses were conducted using data from the search period, including multiple fixations on a distractor or on a target, prior to the final fixation on the target lasting longer than 800 ms. We evaluated the effects of feature-based attention in foveal units by comparing their responses to the same stimulus when it served as a target versus when it served as a distractor, with all fixations followed by saccades. Because spatial attention was directed to the same stimulus in both conditions, the response differences primarily reflect differences in feature-based attentional states. We then examined spatial attention effects in peripheral units by comparing responses when the animal was preparing saccades toward versus away from the RFs.

We found that feature-based attention enhanced the responses of foveal units to target stimuli in both V4 and IT (see **Fig. 2A, B** for the normalized population firing rates averaged across face-selective foveal units during fixations on face targets, face distractors, house targets, and house distractors in V4 and IT, respectively). Responses to face stimuli were significantly higher when they were targets compared to when they were distractors (Wilcoxon signed-rank test, *P* < 0.05). These face-selective units also showed weakly but significantly enhanced responses to house targets in IT (*P* < 0.05), but not in V4 (*P* = 0.89). For house-selective foveal units in both V4 and IT, feature-based attention significantly enhanced responses to house targets (**Fig. 2C, D**). These house-selective units showed small attentional effects in response to face stimuli (*P* < 0.05). Overall, our findings suggest that feature-based attention selectively enhances responses to stimuli in the preferred category of foveal category-selective units in both V4 and IT. For units that exhibited no clear preference for either face or house stimuli, responses to face and house targets were significantly higher than responses to the same stimuli when they served as distractors (Wilcoxon signed-rank test, *P* < 0.05; **Fig. 2E, F**; see also the responses to faces and houses in V4 and IT, respectively, in **Fig. S2G, I**).

**Fig. 2.**
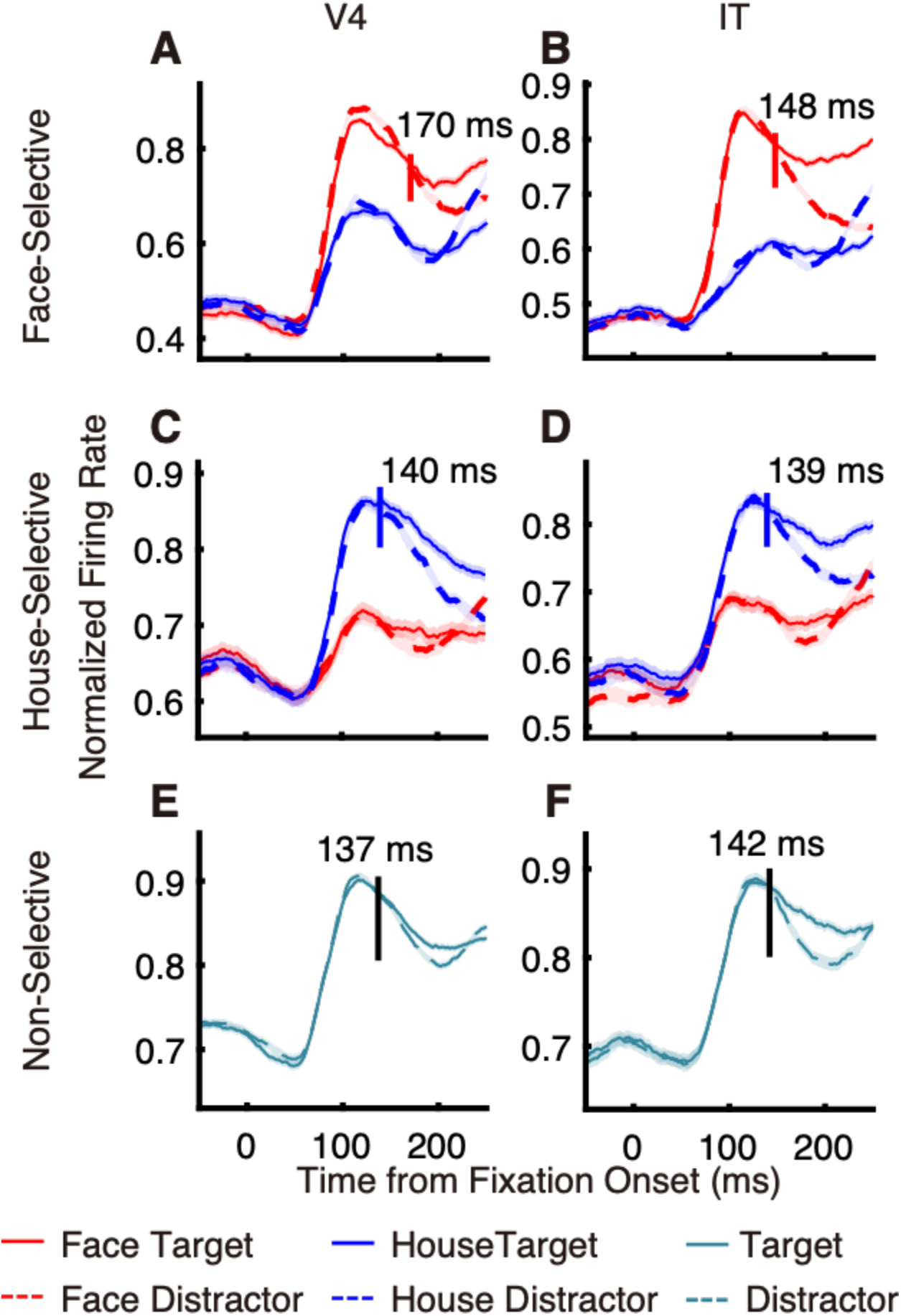
Foveal feature-based attentional modulation in V4 and IT. **(A)** V4 face-selective units. **(B)** IT face-selective units. **(C)** V4 house-selective units. **(D)** IT house-selective units. **(E)** V4 non-selective units. **(F)** IT non-selective units. Shown are normalized population firing rates during fixations on face targets, face distractors, house targets, and house distractors, respectively (see **Methods**). Shading around the mean firing rates indicates ±SEM across units. Vertical lines and numbers indicate population-level latencies. For selective units, firing rates were normalized to the maximum response to the attended stimulus from the preferred category. For non-selective units, fixations on face and house targets were combined into target fixations, and fixations on face and house distractors were combined into distractor fixations. Firing rates were normalized to the maximum response during target fixations.

To further quantify feature-based attention effects, we calculated a feature attention index, defined as the difference between firing rates in the two attention conditions (target versus distractor), normalized by their sum (see **Fig. S2A, B** for the distributions of feature attention index for all foveal face-selective units in V4 and IT, based on their responses to face stimuli). Notably, the average feature attention index was significantly greater than zero (V4: 0.023 ± 0.0049; IT: 0.058 ± 0.0052; Wilcoxon signed-rank test, both *P*s < 0.05). Similarly, the feature attention index of foveal house-selective units in V4 and IT, based on responses to house stimuli, was also significantly greater than zero (**Fig. S2C, D**; V4: 0.030 ± 0.0050, *P* < 0.05; IT: 0.030 ± 0.0062, *P* < 0.05). In both V4 and IT, the feature attention indices calculated from responses to the preferred category were significantly higher than those calculated from responses to the non-preferred category for both face- and house-selective units (Wilcoxon signed-rank test, both *P*s < 0.05). Furthermore, for non-selective units, the feature attention indices calculated from responses to both face and house stimuli were also significantly greater than zero (**Fig. S2E, F**; V4: 0.0091 ± 0.0016, *P* < 0.05; IT: 0.026 ± 0.0043, *P* < 0.05; see also the correlation between feature attention index of faces and houses in V4 and IT, respectively, in **Fig. S2H, J**). We also obtained similar results when controlling for fixation durations on targets and distractors (i.e., there was no significant difference between fixation durations on targets and distractors; Wilcoxon signed-rank test, P > 0.05; **Fig. S2K–P**).

We next analyzed the onset latency of feature attention effects at the *population* level (**Fig. 2**; see **Methods**). For foveal face-selective units, response modulation by feature attention became significant (Wilcoxon signed-rank test, *P* < 0.05) at 170 ms and 148 ms after fixation onset in V4 and IT, respectively. Notably, the attentional effects emerged significantly earlier in IT than in V4 (two-sided permutation test, *P* < 0.05). In contrast, the attentional latencies for house-selective units (V4: 140 ms; IT: 139 ms) and non-selective units (V4: 137 ms; IT: 142 ms) were not significantly different between the two areas. We also computed attentional latency for *individual* foveal units and derived similar results (**Fig. 3A**). Together, we observed a significantly earlier attentional effect in IT than in V4 for face-selective units, but not for house-selective or non-selective units.

**Fig. 3.**
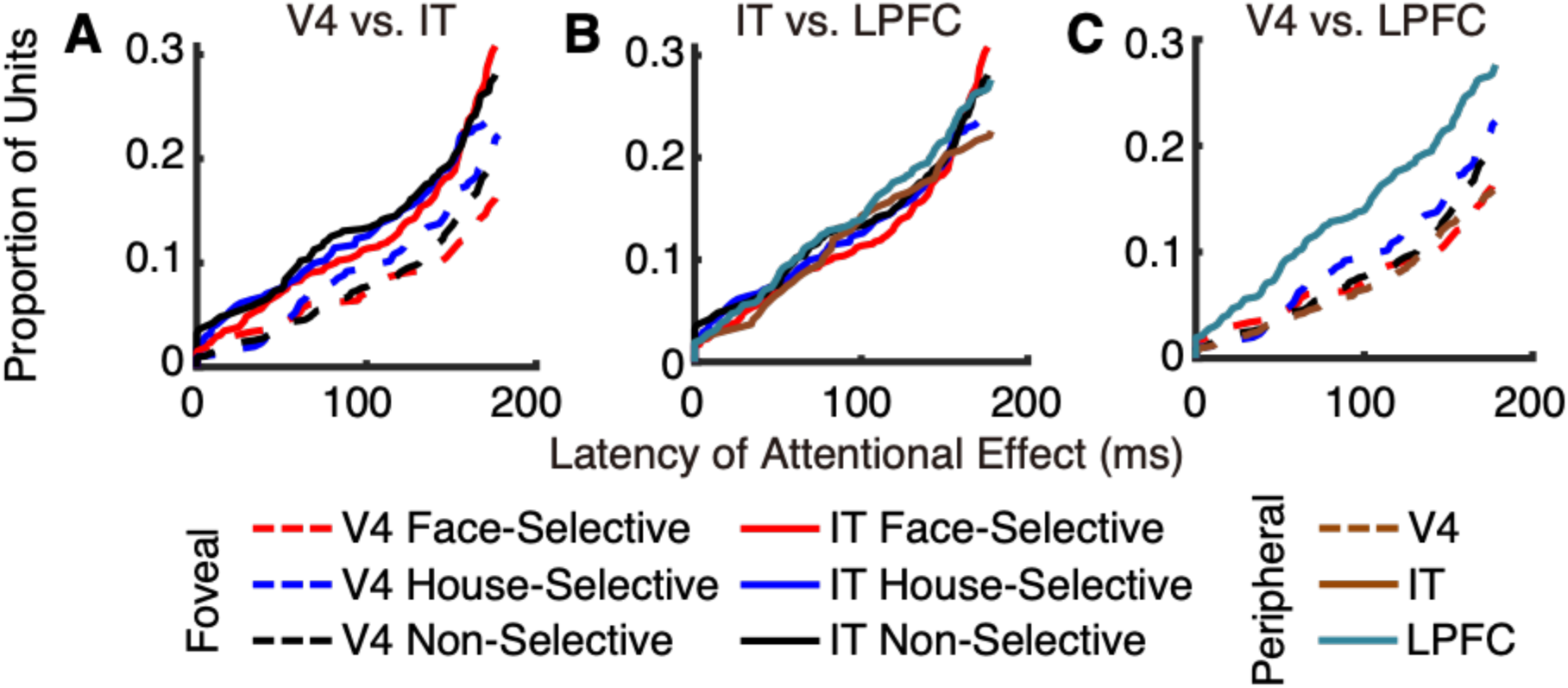
Response latency of attentional modulation in foveal and peripheral receptive fields. Shown are cumulative distributions of feature-attention effect latencies, computed from individual foveal face-, house-, and non-selective units in V4 and IT, and from peripheral non-selective units in V4, IT, and LPFC. **(A)** V4 versus IT. **(B)** IT versus LPFC. **(C)** V4 versus LPFC.

### Feature and spatial attentional modulation of foveal and peripheral units

Of the 765 peripheral units in V4, there were 19 face-selective units, 13 house-selective units, and 730 non-selective units. In IT, the 239 peripheral units included 11 face-selective units, 10 house-selective units, and 216 non-selective units. Among the 507 peripheral units in LPFC, there were 15 face-selective units, 23 house-selective units, and 466 non-selective units (see **Fig. S1L, P, T** for the distributions of selectivity index for all peripheral units). Given the limited number of selective units, we focused our analysis on the non-selective units in V4, IT, and LPFC.

To assess the effects of feature-based attention on units with peripheral RFs without the confounding influence of spatial attention, we examined neural responses when spatial attention was directed outside the RF. Specifically, we compared responses to the same stimuli in the peripheral RFs under two conditions: when the stimuli served as targets and when they served as distractors (see **Methods**). These peripheral feature-based attention effects on visual responses were significant in V4, IT, and LPFC (Wilcoxon signed-rank test, *P* < 0.05; **Fig. 4A–C**), when the monkeys were planning an eye movement toward a stimulus located outside the RF. Similar response patterns were observed when neural activity was aligned to saccade onset (**Fig. S3A–C**).

**Fig. 4.**
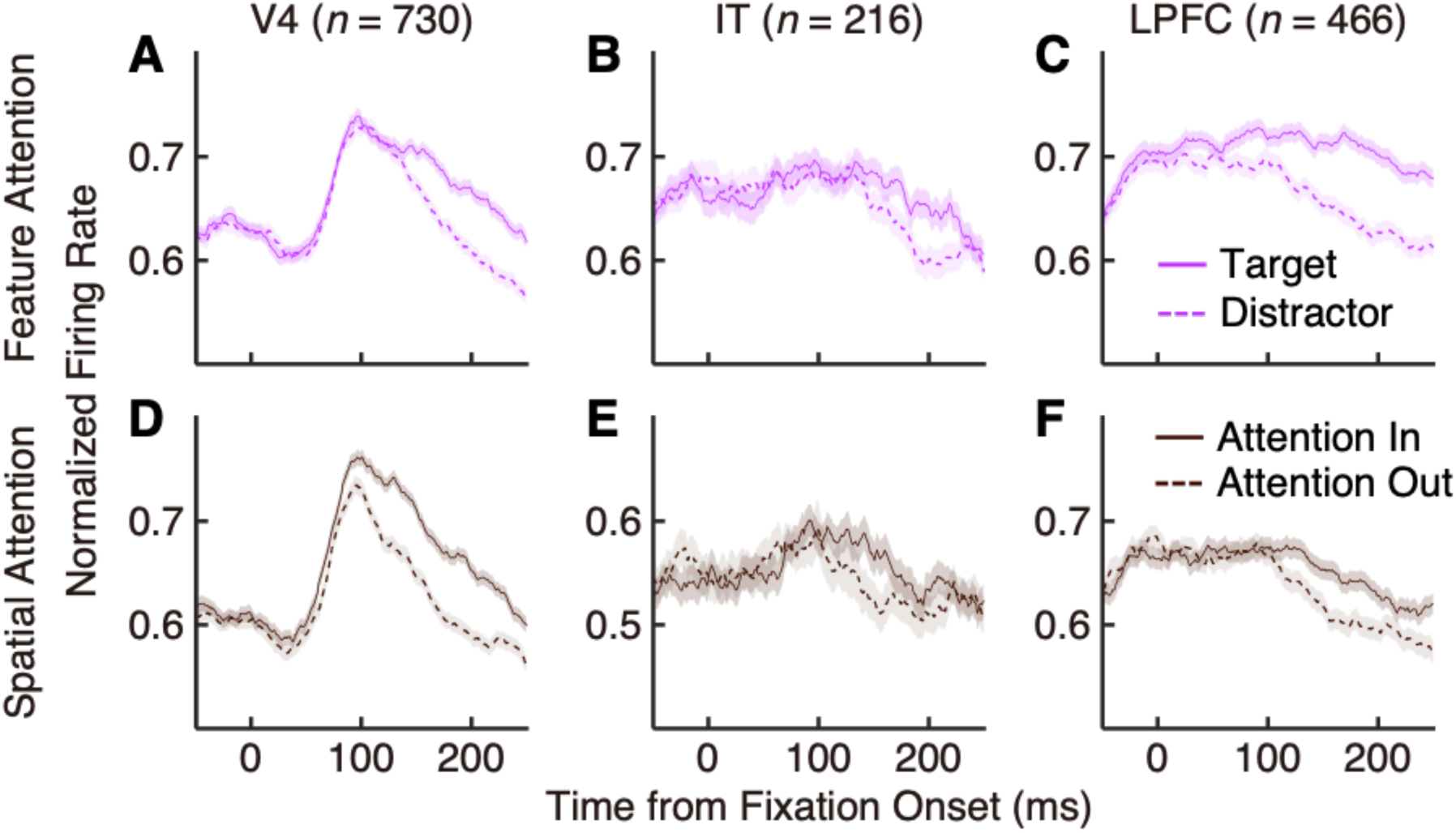
Peripheral feature and spatial attentional modulation in V4, IT, and LPFC. **(A–C)** Population responses of peripheral non-selective units in V4, IT, and LPFC, respectively, to target stimuli and to the same stimuli when they appeared as distractors. **(D–F)** Population responses of the same units to stimuli followed by saccades directed into their receptive fields (attention in) versus away from their receptive fields (attention out), in V4, IT, and LPFC, respectively. **(A, D)** V4. **(B, E)** IT. **(C, F)** LPFC. Shaded area denotes ±SEM across units.

We also analyzed the onset latency of feature-based attention effects in peripheral units. The latencies of these effects were 137 ms, 147 ms, and 58 ms in V4, IT, and LPFC, respectively. The attentional effect in LPFC emerged significantly earlier than in V4 across both foveal and peripheral units (**Fig. 3C**; two-sided permutation test, *P*s < 0.05), consistent with previous findings (5, 6). However, this was not the case for IT (**Fig. 3B**; all *P*s > 0.05). We further compared the latencies of attentional modulation between peripheral and foveal units within the same brain areas. Overall, the latencies of feature attention effects did not differ significantly between foveal and peripheral units in V4 and IT (all *P*s > 0.05; **Fig. 3B, C**).

To assess the effects of spatial attention on units with peripheral RFs, we compared neural responses to the same stimulus in the RF under two conditions: when the animal was preparing a saccade to that stimulus (“attention in”) and when the animal was preparing a saccade away from the RF (“attention out”). We found that attention-in responses were significantly greater than attention-out responses in V4, IT, and LPFC (Wilcoxon signed-rank test, *P* < 0.05; **Fig. 4D–F**) and immediately before saccade onset (Wilcoxon signed-rank test, *P* < 0.05; **Fig. S3D–F**).

### Distribution of feature and spatial attention in the entire visual field during search

To examine feature and spatial attention across the visual field during search, all fixations prior to the final target fixation were categorized into four types: (1) “Distractor fixations to D”, during which the monkey fixated on a distractor and prepared a saccade to another peripheral distractor; (2) “Distractor fixations to T”, during which the monkey fixated on a distractor and prepared a saccade to a target; (3) “Target fixations to D”, during which the monkey fixated on a target and prepared a saccade to a distractor; and (4) “Target fixations to T”, during which the monkey fixated on a target and prepared a saccade to the other target. The features of the stimulus in the foveal field were not attended during the two types of distractor fixations, but were attended during the two types of target fixations.

For units with peripheral RFs across V4, IT, and LPFC, during both “Distractor fixations to D” and “Distractor fixations to T”, feature attention enhanced responses to targets in the periphery, and spatial attention enhanced responses to the saccade target (Wilcoxon signed-rank test, *P* < 0.05; **Fig. 5A–F**). These significant feature- and spatial-attention enhancements remained evident when neural activity was aligned to saccade onset (**Fig. S4A–F**). Notably, distinct response patterns for feature attention emerged when the animals were fixating on a target. In contrast to distractor fixations, during “Target fixations to D”, there was no significant feature attentional enhancement in response to the peripheral target (Wilcoxon signed-rank test, *P* > 0.05; **Fig. 5G–I** left panels). That is, during these fixations, it made no difference whether the stimulus in the peripheral RF was a target or a distractor. Similar response patterns were observed when neural activity was aligned to saccade onset (**Fig. S4G–I** left panels). During “Target fixations to T”, because the only peripheral target stimulus was both within the RF and the target of the upcoming saccade, it was not possible to isolate the effects of feature attention independent of spatial attention. Therefore, we did not assess feature attentional effects under this condition. We further found that spatial attentional enhancements to the saccade target were reduced during target fixations compared to distractor fixations in V4 and IT when activity was aligned to fixation onset (Wilcoxon rank-sum test, *P* < 0.05; **Fig. 5G, H** versus **Fig. 5A, B**), although this effect was not completely abolished. However, spatial attention effects during target fixations remained evident when aligned to saccade onset (**Fig. S4G–I**).

**Fig. 5.**
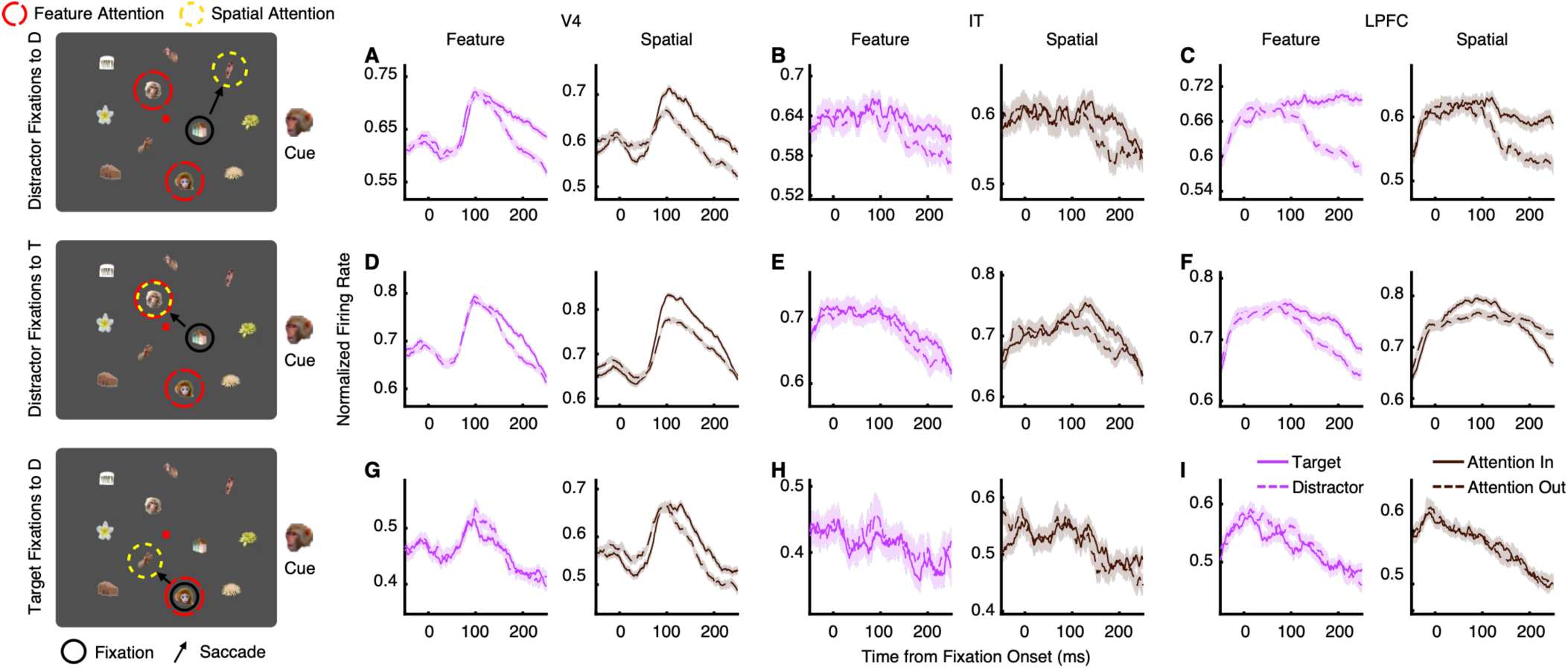
Feature and spatial attention in peripheral non-selective units. Left panels show average population responses to target stimuli and distractor stimuli (i.e., feature attention). Right panels show responses of the same units to stimuli followed by saccades into their RFs (attention in) or out of their RFs (attention out) (i.e., spatial attention). **(A–C)** The current fixation is on a distractor and the subsequent fixation is also on a distractor. **(D–F)** The current fixation is on a distractor and the subsequent fixation is on a target. **(G–I)** The current fixation is on a target and the subsequent saccade is on a distractor. **(A, D, G)** V4 (*n* = 727). **(B, E, H)** IT (*n* = 216). **(C, F, I)** LPFC (*n* = 463).

Together, by integrating foveal and peripheral attentional effects across different fixation types, our study reveals systematic patterns of feature and spatial attention across the visual field during visual search.

### Impact of feature-based attention on saccade behavior

The distribution of feature and spatial attention had an important impact on subsequent saccade behaviors throughout the search period. On average, target fixations (256.69 ± 197.44 ms [mean ± SD]) were significantly longer than distractor fixations (156.26 ± 45.94 ms; Wilcoxon rank-sum test, *P* < 0.0001), and during these prolonged target fixation, foveal feature-based attention modulation was consistently observed. Saccades following distractor fixations could be guided by both peripheral feature and spatial attention, whereas peripheral feature attention was absent during target fixations. Consistently, the probability of making a saccade to a peripheral target was higher following distractor fixations (75.22%) than following target fixations (48.44%, or 63.49% after probability calibration; see **Methods**), indicating the important role of peripheral feature-based attention in guiding eye movements. However, the relative probability of saccades to targets following target fixations—normalized to the number of target and distractor stimuli within the search array—was still higher than that of saccades to distractors. This suggests that additional mechanisms may contribute to guiding saccades to peripheral targets following target fixations.

We analyzed four types of saccade sequences surrounding current target fixations (T), based on the type of fixations (target or distractor) immediately before and after the target fixation (**Fig. 6A**). Among them, T_pre_→T→T_pre_ and D_pre_→T→T_post_ were the two sequences in which saccades were directed toward a target stimulus following the target fixation. In the T_pre_→T→T_pre_ sequence, animals made saccades back to the previously fixated target stimulus (T_pre_) after the current target fixation. Consistently, the response enhancement during T_pre_ fixations was stronger for targets that would be re-fixated compared to those that would not, in both V4 and IT foveal units (Wilcoxon signed-rank test, *P* < 0.05; **Fig. 6B–G**). The activity was higher around saccades directed to the previously fixated target than around saccades directed to a distractor in both V4 and IT foveal units (Wilcoxon signed-rank test, *P* < 0.05; **Fig. S5**). In the D_pre_→T→T_post_ sequence, we observed stronger enhancement for peripheral targets (T_post_)—which would become the targets of the second following saccade—in peripheral units in V4, IT, and LPFC during the D_pre_ fixations (**Fig. 6H–J**). Together, feature-based attention effects during these T_pre_ and D_pre_ fixations may contribute to the selection of saccade targets following target fixations. This may help explain the relatively high probability of saccades to targets following target fixations, even in the absence of peripheral feature attention effects.

**Fig. 6.**
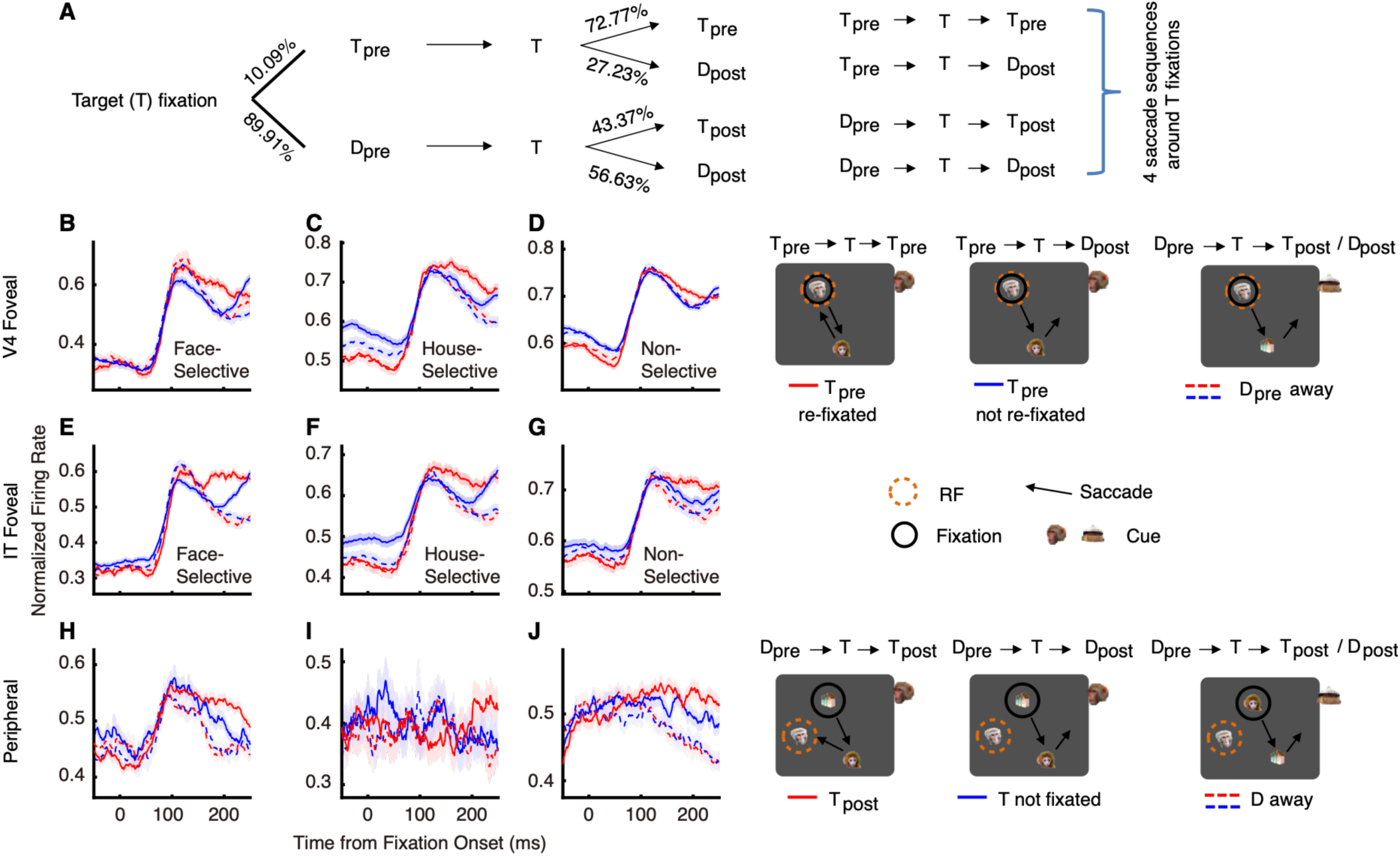
Feature attentional modulation and saccade transitions around target fixations. **(A)** Four saccade sequences around target (T) fixations. T_pre_ and D_pre_ represent the target and distractor fixations immediately before the target fixation, respectively. T_post_ and D_post_ represent the target and distractor fixations immediately after the target fixation, respectively. Arrows (→) indicate saccades. In the T_pre_**→**T**→**T_pre_ sequence, the pre- and post-target fixations were on the same stimulus. The proportions of trials/saccades for each sequence are shown along the lines/arrows (left panel). A summary of the four saccade sequences is shown in the right panel. **(B–G)** Average population responses to the fixated target stimulus in the T_pre_ fixation of the T_pre_→T→T_pre_ (‘T_pre_ re-fixated’, red solid line) and T_pre_→T→D_post_ (‘T_pre_ not re-fixated’, blue solid line) sequences, and to the same distractor stimulus in the D_pre_ fixation of the D_pre_→T→T_post_/D_post_ sequences (the red dashed line indicates ‘D_pre_ away’ fixations matched with ‘T_pre_ re-fixated’ fixations; the blue dashed line indicates ‘D_pre_ away’ fixations matched with ‘T_pre_ not re-fixated’ fixations). **(B)** IT foveal face-selective units (*n* = 476). **(C)** IT foveal house-selective units (*n* = 336). **(D)** IT foveal non-selective units (*n* = 556). **(E)** V4 foveal face-selective units (*n* = 246). **(F)** V4 foveal house-selective units (*n* = 304). **(G)** V4 foveal non-selective units (*n* = 1051). The corresponding behavioral conditions are illustrated on the right side of the plots. Shading around the mean firing rates indicates ±SEM. **(H–J)** Population responses of peripheral non-selective units to target stimuli in their receptive fields in D_pre_ fixations of the D_pre_→T→T_post_ (‘T_post_’, red solid line) and D_pre_→T→D_post_ (’T not fixated’, blue solid line) sequences, and to the same distractor stimulus in the D_pre_ fixation of the D_pre_→T→T_post_/D_post_ sequences (the red dashed line indicates ‘D_pre_ away’ fixations matched with ‘T_pre_ re-fixated’ fixations; the blue dashed line indicates ‘D_pre_ away’ fixations matched with ‘T_pre_ not re-fixated’ fixations). **(H)** V4 (*n* = 566). **(I)** IT (*n* = 103). **(J)** LPFC (*n* = 414). Behavioral conditions for the peripheral units are illustrated on the right side of the plots.

### Dependence of feature attentional modulation on stimulus category

While feature attention to simple visual features—such as color, shape, luminance, and motion direction—has been widely studied, there remains a lack of research on feature attention to more complex attributes, such as object categories in naturalistic images. In this study, we observed greater attentional enhancement for stimuli in the preferred category in foveal units (**Fig. 2**), along with a wide range of responses to different stimuli within the same category. To further clarify the role of category features in attentional modulation, we analyzed attentional effects on responses to different stimuli within the same category and across different categories. Stimuli from the same category were divided into four subsets based on their response amplitudes (see **Fig. 7A–H** for illustration of the attentional effects across subsets of house and face stimuli in IT house-selective units, ordered from low to high response amplitudes; see **Fig. 7I** for the average attentional effects [150–225 ms after fixation onset] across these subsets). The visual response amplitudes to the four subsets of house stimuli differed significantly both when they were attended (one-way ANOVA, F(3,335) = 20.33, *P* < 0.001; **Fig. 7A–D** solid line) and unattended (F(3,335) = 19.57, *P* < 0.001; **Fig. 7A–D** dashed line; **Fig. 7I**). However, the attentional enhancement across the four subsets did not differ significantly (one-way ANOVA, F(3,335) = 1.09, *P* > 0.05). In contrast, the attention effects for different categories (house versus face) were significantly different (one-way ANOVA, F(1,1354) = 19.09, *P* < 0.001). These results suggest that feature-based attentional modulation may depend more on the categorical identity of the stimulus than on the magnitude of the visual response it elicits.

**Fig. 7.**
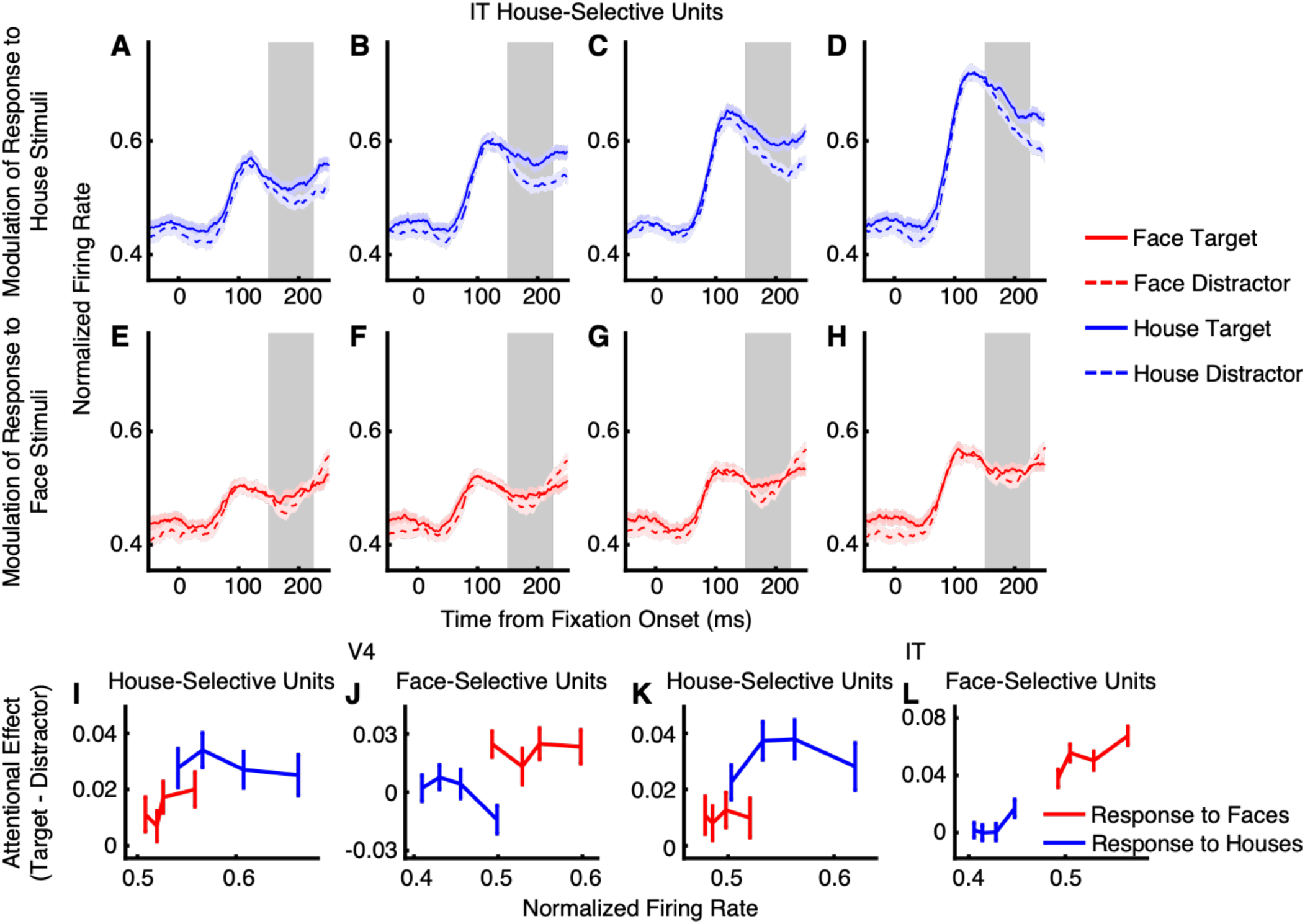
Influence of stimulus category on feature attentional modulation. **(A–D)** Feature attention effects on responses to four subsets of house stimuli for IT house-selective units (*n* = 339). The subsets were sorted from low to high based on the amplitude of the responses to house distractor stimuli. The shaded areas indicate the time window (150–225 ms after fixation onset) used for analyzing attentional effects. **(E–H)** Feature attention effects on responses to four subsets of face stimuli for the same IT house-selective units (*n* = 339). **(I–L)** Attentional effects as a function of visual response amplitudes to subsets of face and house stimuli. **(I)** IT house-selective units (*n* = 339). **(J)** IT foveal face-selective units (*n* = 480). **(K)** V4 foveal house-selective units (*n* = 301). **(L)** V4 foveal face-selective units (*n* = 266). Normalized visual response amplitudes to subsets of house and face distractors (calculated in a window from 50–225 ms after fixation onset) are plotted against the amplitude of attentional effects (attended – unattended, calculated in a window from 150–225 ms after fixation onset).

These tendencies were also observed in V4 house-selective units (within the preferred category: one-way ANOVA, F(3,297) = 0.33, *P* > 0.05; across categories: F(1,1202) = 10.73, *P* < 0.001; **Fig. 7K**; **Fig. S6I–** and V4 face-selective units (within the preferred category: F(3,362) = 0.46, *P* > 0.05; across categories: F(1,1062) = 15.97, *P* < 0.001; **Fig. 7L**; **Fig. S6Q–X**). For IT face-selective units, the attentional effects were significantly different both within the preferred category and across categories (one-way ANOVA, *P* < 0.05; **Fig. 7J**; **Fig. S6A–H**), but the effect size across categories (F(1,1918) = 125.65) was substantially larger than that within the preferred category (F(3,476) = 3.77).

## Discussion

In this study, we investigated attentional processing in both the foveal and peripheral visual fields during a free-gaze visual search task. In this task, attention is strongly guided by top-down goals, which bias processing toward behaviorally relevant features and object categories (2, 50, 51). Top-down attention, including categorical and feature-specific components, has been shown to modulate neural processing across the visual pathway based on task demands and to originate from distributed frontoparietal control networks (11, 35–38, 40). Our study provides further insight into the mechanisms of goal-directed visual attention, as it is among the first to demonstrate foveal feature attention effects during free-gaze visual search, as well as the distribution of feature and spatial attention across the entire visual field. The search period included multiple fixations on both targets and distractors before the final fixation on the target. During target fixations, enhanced responses to the foveal target stimuli—reflecting feature-based attention—were associated with longer fixation durations (compared to distractor fixations on the same stimuli), a higher likelihood of subsequent re-fixation on the currently fixated target, and a reduction or delay in both feature- and spatial-attention effects in the peripheral field. These findings suggest that foveal attentional enhancement supports maintaining gaze on the current target or returning to it later. In contrast, during distractor fixations, feature-based attention in the periphery was associated with a higher probability of saccades toward peripheral target stimuli, typically occurring within one or two eye movements following the current fixation. Together, these results indicate a complementary interaction between foveal and peripheral feature-based attentional processes in guiding efficient visual search, highlighting the critical role of foveal feature attention in active vision.

Consistent with the primate visual system’s preferential processing of foveal stimuli, we recorded a substantial proportion of units with small RFs covering the foveal region in V4 (40.27%) and IT (30.28%). These units exhibited stronger category selectivity than those with peripheral RFs. Approximately 57% of foveal units in IT and 30% in V4 were selective for faces or houses, whereas only 8% of peripheral units in IT and 4% in V4 showed category selectivity. This bias may support the visual processing of complex stimuli in the foveal field. Moreover, we observed feature-based attentional enhancements in foveal units responding to target stimuli in both V4 and IT. In contrast, prior studies of feature attention have primarily focused on units with peripheral RFs (3, 5, 6, 8, 9, 52–55). The category-selective foveal units showed stronger attentional enhancements for stimuli in their preferred category, whereas non-selective units exhibited similar enhancements across all categories. Together, these findings extend our understanding of the spatial distribution of feature attention effects across the entire visual field during active visual search.

Regarding attentional processing in the foveal and peripheral visual fields, our results suggest that foveal feature-based attention may outweigh peripheral attentional processing in the context of our task. Specifically, peripheral feature-based attentional enhancements disappeared when attention was directed to the feature of a foveal stimulus. In contrast, foveal attention effects persisted even when a peripheral target—capable of eliciting feature-based enhancements in certain peripheral units—was present. These findings challenge the commonly held view that feature-based attention is uniformly distributed across the entire visual field (3, 5, 6, 8, 9, 13–15, 56, 57). Instead, they suggest that global feature attention effects observed in previous studies may be largely confined to the peripheral visual field. Furthermore, foveal feature-based attention may modulate peripheral spatial attention effects in both V4 and IT. By integrating these attentional effects across foveal and peripheral fields during different types of fixations throughout the search process, our study reveals novel and comprehensive patterns of feature and spatial attention distribution across the entire visual field.

Some studies have provided evidence for integration between peripheral and foveal feature information across saccades, including features such as stimulus color (58, 59) and object orientation (60, 61), and visual features have been shown to be predictively remapped prior to saccades (62). Our finding provides a potential neuronal mechanism that may support this integration process (63). We found that LPFC’s extensive representation of the visual periphery provides a neural substrate for monitoring the broader search array. Crucially, our finding that LPFC activity temporally precedes attentional effects in the visual area—consistent with previous studies (6, 9, 11, 35–40)—suggests that it does not merely reflect peripheral sensory input. Instead, LPFC likely acts as a top-down orchestrator, projecting task-relevant templates derived from current foveal goals onto peripheral candidate locations, a possibility that warrants further investigation.

The distribution of attentional effects is crucial for guiding saccades during efficient visual search. Models such as Guided Search and the priority map propose a collaboration between parallel and serial processing: parallel processing evaluates multiple stimuli across the visual field to generate a priority map that integrates stimulus salience with top-down factors, including feature-based attention; serial processing then directs eye movements to high-priority locations on this map (2, 41, 50). Our findings show that parallel feature-based attention effects in the peripheral field depend on the attentional state at the fovea, which itself results from serial selection. This suggests that parallel and serial attention processes operate within a bi-directionally interactive loop, working together to support effective visual search.

We used naturalistic, complex stimuli in our visual task, closely resembling the objects encountered in daily life. Feature-based attention to simple visual features—such as color, shape, or motion—has been extensively studied (1, 3, 5, 7–9, 11, 12, 64). Attention can also operate over more complex features. For example, objects themselves can serve as units of attentional selection (65, 66), and feature-based attentional effects have been observed when searching for images that match the cued images or image patches (6, 67). In this context, attention can be directed either to overall features of an object or to objects as configurations of multiple non-spatial features. Furthermore, attention to the category of stimuli has been extensively investigated in fMRI experiments in humans (68–71), and it has been shown that attention can warp the representations of semantically related categories when participants search for different categories (70). In this study, the neural responses to targets versus distractors were compared while constrained to the same stimuli across different trials, ensuring that the observed response divergence was not due to the physical category of the targets and distractors. We found that face-selective and house-selective units exhibited stronger feature-based attentional enhancements for stimuli in their preferred category, similar to selective attention effects observed for simple features in previous research (1, 3, 72). Moreover, the attentional effects were consistent across stimuli within the same category, regardless of differences in response magnitude. This suggests that feature-based attention was driven by stimulus category rather than response level alone. Thus, similar to attention guided by simple features, feature-based attention can also operate on complex features in naturalistic stimuli.

A major challenge in interpreting neural activity related to attentional modulation is the inherent temporal overlap of visual processing, motor planning, and target recognition signals in the free-gaze visual search task (73). To isolate genuine feature-based attention from potential confounds, we applied several stringent analytical constraints, consistent with prior studies (3, 5, 6). Specifically, by restricting our analysis to fixations where the subsequent saccade was directed away from the RFs, we dissociated attentional modulation from the preparatory motor activity associated with saccade execution. Furthermore, by comparing responses to the same physical stimulus—alternating its role as a target or distractor across trials—we eliminated any potential bias introduced by stimulus identity or physical category. We restricted our analysis to fixations preceding target selection—that is, before the monkeys made a behavioral choice—to minimize contributions from target detection or decision-related signals.

An important caveat is that small eye movements, such as microsaccades or fixation instabilities, may disproportionately influence responses in foveal compared to peripheral RFs, given the smaller size of foveal receptive fields. Although this concern is particularly pronounced in early visual areas such as V1, it is likely less severe in higher-level areas such as IT, where RFs are substantially larger and more tolerant to slight gaze shifts. Nevertheless, we cannot fully exclude the possibility that such eye movements contributed to variability in our foveal responses. Future work incorporating higher-resolution eye tracking and analyses specifically designed to quantify microsaccade-related effects will be important for determining the extent to which subtle eye movements shape foveal activity during naturalistic free viewing.

## Acknowledgments

We thank Robert Desimone and Shuo Wang for their valuable comments. This research was supported by the National Natural Science Foundation of China (62027804, 62206141, 31671108, and 62236009), International Partnership Program of Chinese Academy of Sciences (172644KYSB20160175), Shenzhen Fundamental Research Program (JCYJ20200109114805984), and the major key project of the Peng Cheng Laboratory. The funders had no role in study design, data collection and analysis, decision to publish, or preparation of the manuscript.

## Author Contributions

Conceptualization, H.Z.; Methodology, H.Z., R.D., J.Z., Z.M., X.Z., and H.E.; Software, J.Z. and H.Z.; Investigation, J.Z. and X.Z.; Formal Analysis, J.Z., Z.M., and X.Z.; Resources, H.Z. and Y.T.; Writing – Original Draft, H.Z., Z.M., and J.Z.; Writing – Review & Editing, H.Z., Shuo Wang, Z.M., R.D., Y.T., Y.W., and Shanshan Wang; Funding Acquisition, H.Z., Z.M., and Y.T.; Supervision, H.Z., R.D., and Z.M.; Project Administration, H.Z.

## Methods

### Subjects and recording sites

Two male rhesus macaques, weighing 12 and 15 kg, were used in this study. Under aseptic conditions, the monkeys were implanted with a post to stabilize the head and recording chambers over areas V4, IT, and LPFC. The chamber locations were determined based on MRI scans obtained prior to surgery. Recordings in IT spanned the central IT cortex, encompassing the area between the anterior middle temporal sulcus (AMTS) and the posterior middle temporal sulcus (PMTS), including TE and TEO. Recordings in LPFC were located anterior to the arcuate sulcus (AS) and lateral to the principal sulcus (PS), mainly covering areas 45 and 44. All experiments were conducted at the Shenzhen Institutes of Advanced Technology, Chinese Academy of Sciences, with approval from the Institutional Animal Care and Use Committee (No. SIAT-IRB-160223-NS-ZHH-A0187-003). This dataset has been analyzed in previous studies (44–46).

### Tasks and stimuli

Monkeys were trained to perform a free-gaze visual search task. A central fixation point was presented for 400 ms, followed by a cue lasting 500 to 1300 ms. After a delay of 500 ms, the search array appeared. The search array contained 11 items, including two targets, which were randomly selected from a total of 20 predefined locations. Monkeys were required to find one of the two targets within 4000 ms and maintain fixation on the target for 800 ms to receive a juice reward. No constraints were placed on their search behavior, allowing the animals to perform the search naturally. Before the onset of the search array, monkeys were required to maintain central fixation. The two target stimuli belonged to the same category as the cue stimulus, though they were distinct images. We utilized four categories of stimuli—face, house, flower, and hand—each comprising 40 images. The cue stimulus was randomly selected from either the house or face category with equal probability. The remaining 9 stimuli in the search array were drawn from the other three categories. Each stimulus subtended an area of approximately 2° × 2°, with hue, saturation in the HSV color space, aspect ratio, and luminance matched across categories (44–46). The 20 locations, covering the visual field with eccentricities from 5° to 11°, included 18 locations symmetrically distributed across the left and right visual fields (9 on each side), and 2 locations on the vertical midline.

A visually guided saccade task was used to map the peripheral receptive fields (RFs) of recorded units. After a 400-ms central fixation, a stimulus (a face or house, identical to those in the visual search task) randomly appeared at one of 20 locations. Monkeys were required to make a saccade to the stimulus within 500 ms and maintain fixation on it for 300 ms to receive a reward.

Behavioral experiments were conducted using MonkeyLogic software (University of Chicago, IL), which presented stimuli, monitored eye movements, and triggered reward delivery.

### Electrophysiology

Single-unit and multi-unit spikes were recorded from V4, IT, and LPFC using 24- or 32-contact electrodes (V-Probe or S-Probe, Plexon Inc., Dallas, USA) with a 128-channel Cerebus System (Blackrock Microsystems, Salt Lake City, UT, USA). In most sessions, activity was recorded simultaneously from two of these areas. Neural recordings were filtered between 250 Hz and 5 kHz and digitized at 30 kHz to obtain spike data. Spike sorting was performed using Plexon’s Offline Sorter™ (OFS). LFP signals were obtained by filtering neural recordings between 0.3 and 250 Hz and digitizing them at 1000 Hz. Recording locations in V4, IT, and LPFC were verified using MRI. Eye movements were recorded with an infrared eye-tracking system (iViewX Hi-Speed, SensoMotoric Instruments [SMI], Teltow, Germany) at a sampling rate of 500 Hz.

### Data analysis: spike rate

Measurements of neural activity were obtained from spike density functions, which were generated by convolving the time of action potentials with a function that projects activity forward in time (Growth = 1 ms, Decay = 20 ms) and approximates an EPSP (74). Specifically, this spike density function has two advantages (74). First, each spike exerts influence only forward in time, representing the actual postsynaptic effect of each cell. Second, by using a function that resembles a postsynaptic potential, we can apply time constants similar to those measured physiologically. Additionally, using the postsynaptic potential filter for time course analysis is advantageous because, when using the Gaussian filter, target discrimination times sometimes occur earlier than the unit’s evident visual latency. This impossible outcome occurs because, with the Gaussian filter, spikes exert influence backward in time.

### Data analysis: receptive field

The visual response to the cue and the search array in the free-gaze visual search task was assessed by comparing the firing rate during the post-stimulus period (50 to 200 ms after cue/array onset) to the corresponding baseline (−150 to 0 ms relative to cue/array onset) using a Wilcoxon rank-sum test. Based on these responses, we classified units into three categories of RFs:

i. Units with a focal foveal RF: These units responded solely to the cue in the foveal region (*P* < 0.05) but not to the search array that included items in the periphery (*P* > 0.05).
ii. Units with a broad foveal RF: These units responded to both the cue and the search array (both *P*s < 0.05).
iii. Units with a peripheral RF: These units only responded to the search array (*P* < 0.05) but not to the cue (*P* > 0.05). The RFs of these units were additionally mapped based on their activities in the visually guided saccade task. Units whose RFs could be mapped in this task had a localized peripheral RF, whereas units whose RFs could not be mapped had an unlocalized peripheral RF (i.e., units that responded to the search array onset but not during the saccade task; 11.87% of visually responsive units in V4, 5.58% in IT, and 58.29% in LPFC had an unlocalized RF).

Units not classified into the above categories (both Ps > 0.05) were not visually responsive and were excluded from further analysis. In this study, our focus was on units with focal foveal RFs and units with localized peripheral RFs. All further analyses were conducted on these units.

### Data analysis: selection of category-selective units

We selected *category-selective units* by comparing the response to face cues versus house cues in a time window of 50 to 200 ms after cue onset (Wilcoxon rank-sum test, *P* < 0.05). We further imposed a second criterion using a selectivity index similar to indices employed in previous IT studies (75, 76). For each unit with a foveal RF, the response to face stimuli (*R*_face_) or house stimuli (*R*_house_) was calculated using the visual search task by subtracting the mean baseline activity (−150 to 0 ms relative to the onset of the cue) from the mean response to the face or house cue (50 to 200 ms after the onset of the cue). For each unit with a peripheral RF, *R*_face_ and *R*_house_ were calculated using the visually guided saccade task by subtracting the mean baseline activity (−150 to 0 ms relative to the peripheral stimulus onset) from the mean response to the saccade target (50 to 200 ms after the onset of the saccade target). It is worth noting that for both foveal and peripheral units, we ensured that there was only one stimulus in the RF. The selectivity index (SI) was then defined as (*R*_face_ − *R*_house_) / (*R*_face_ + *R*_house_). SI was set to 1 when *R*_face_ > 0 and *R*_house_ < 0, and to −1 when *R*_face_ < 0 and *R*_house_ > 0. Face-selective units were required to have an *R*_face_ at least 130% of *R*_house_ (i.e., the corresponding SI was greater than 0.13). Similarly, house-selective units were required to have an *R*_house_ at least 130% of *R*_face_ (i.e., the corresponding SI was smaller than −0.13). Units were labeled as non-category-selective if the response to face cues versus house cues was not significantly different (*P* > 0.05). The remaining units that did not fit into any of the aforementioned types were classified as undefined units (i.e., there was a significant difference but did not meet the second criterion). It is worth noting that we did not use the activity during the search to calculate the SI to minimize interactions with the attentional effect and between RFs.

### Data analysis: attentional effect

To investigate feature-based attention in foveal units, we compared responses to a stimulus in the fovea when it served as a target versus when the same stimulus served as a distractor, while the monkey was preparing a saccade away from the stimulus. For face- or house-selective units, fixations during the search period (before the final fixation on the target at the end of the search) were sorted into four types: face target, face distractor, house target, and house distractor. In face-target fixations, the stimulus in the fovea was a face, and the monkey was searching for a face target. In face-distractor fixations, the stimulus in the fovea was also a face, but the monkey was searching for a house target. In house-target fixations, the stimulus in the fovea was a house, and the monkey was searching for a house target. In house-distractor fixations, the stimulus in the fovea was a house, and the monkey was searching for a face target. For non-selective units, face-target and house-target fixations were combined into “target” fixations, and face-distractor and house-distractor fixations were combined into “distractor” fixations. The stimulus in the fovea was matched across attended (i.e., target) and unattended (i.e., distractor) conditions. Neural activity in V4 and IT during these fixations was calculated and compared to assess feature attention effects.

To assess feature attention effects in peripheral units, we sorted fixations during the search period, following a procedure similar to that in our previous study (5), into two types: “target” – a target stimulus was located within the unit’s peripheral RF; and “distractor” – the same stimulus appeared in the same peripheral RF location but served as a distractor. Only fixations followed by a saccade away from the RF were included in this analysis. For assessing feature attention effects during “Distractor fixations to D”, “Distractor fixations to T”, and “Target fixations to D”, we analyzed neural responses during fixations followed by saccades away from the RF, thereby excluding the influence of spatial attention. It is important to note that, during “Target fixations to T”, there was no opportunity for the peripheral units to have a target stimulus in their RFs while also being followed by a saccade away from the RF. Therefore, we did not assess feature attention effects for this condition.

To evaluate spatial attention effects in peripheral units, we compared responses between attention-in and attention-out fixations. Attention-in fixations were followed by saccades to a stimulus within the RF, while attention-out fixations were followed by saccades to a stimulus outside the RF. The saccade-target stimulus in the RF during attention-in fixations was matched to a stimulus in the same location during attention-out fixations; in both conditions, this stimulus always served as a distractor for that trial, except in the “Distractor fixations to T” condition (**Fig. 5** and **Fig. S4**), in which it instead served as the target. This design eliminates differences due to feature-based attention between the attention-in and attention-out conditions.

The attention index, used to quantify the magnitude of attention effects, was defined as the difference divided by the sum of firing rates between the two attention conditions, based on the average firing rate within a 150–225 ms window after fixation onset.

### Data analysis: response latency of attentional effects

The latency of attention effects at the population level was determined using the averaged responses of individual units with a sliding window method. A significant difference between target and distractor responses (Wilcoxon signed-rank test, *P* < 0.05) sustained for at least 35 ms was used to define the onset of attentional modulation; specifically, the first time point of that 35 ms window was marked as the onset. To assess whether latency differences at the population level were significant, we conducted a two-sided permutation test with 1,000 runs, as described in our previous study (5).

For single-unit latency estimation, the onset was defined as the first 20 ms bin out of twelve successive bins showing significant differences between the two compared conditions. The latencies of feature attention effects for selective units were calculated based on responses to stimuli in the preferred category.

### Data analysis: probability calibration

In the search array, there were two targets and nine distractors in total. Following distractor fixations, the stimulus array comprised two targets and eight distractors. In contrast, following target fixations, there was only one target and nine distractors. Due to these differences in search space, probability calibration was needed to allow a fair comparison of saccade probabilities toward peripheral targets after different types of fixations. In the condition following distractor fixations, the probability of saccades to peripheral targets was 75.22%, which broke down into 75.22% / 2 = 37.61% to one peripheral target and 24.78% / 8 = 3.0975% to one peripheral distractor. Assuming the same ratio between the probabilities of saccades to peripheral targets versus peripheral distractors held true for both target and distractor fixation conditions, the expected probability of a saccade to a peripheral target following a target fixation was 37.61% / (37.61% + 3.0975% × 9) = 57.42%. However, the observed probability was 48.44%, which, after probability calibration, could be adjusted to 48.44% × 75.22% / 57.42% = 63.49%.

### Data availability

All data that support the findings of this study are publicly available on OSF (https://osf.io/sdgkr/).

### Code availability

The source code for this study is publicly available on OSF (https://osf.io/sdgkr/).

---

**Fig. S1.**
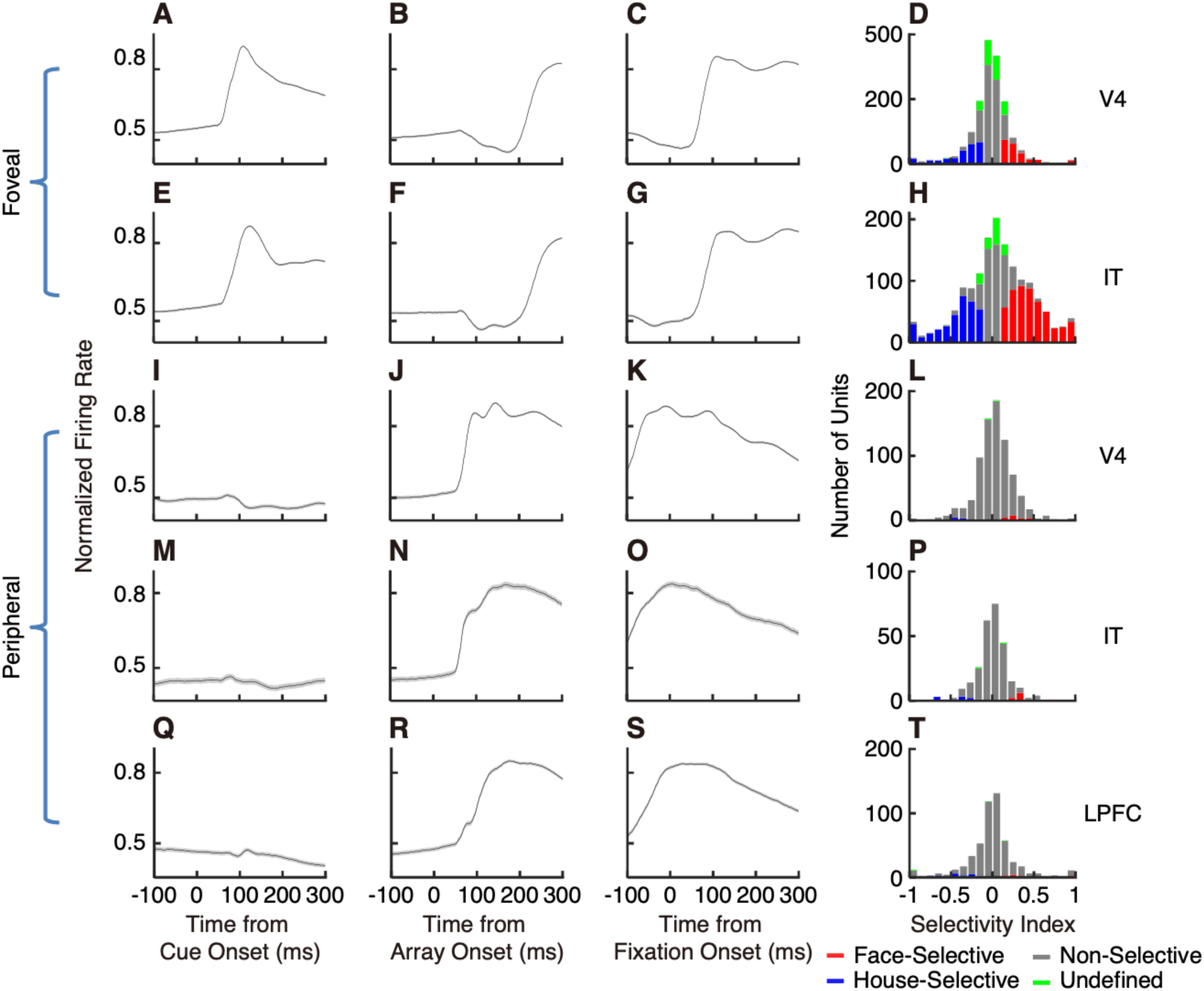
Response of foveal and peripheral units. **(A–D)** V4 foveal units (*n* = 1898). **(E–H)** IT foveal units (*n* = 1511). **(I–L)** V4 peripheral units (*n* = 765). **(M–P)** IT peripheral units (*n* = 239). **(Q–T)** LPFC peripheral units (*n* = 507). **(A, E, I, M, Q)** Mean population response aligned at cue onset. **(B, F, J, N, R)** Mean population response aligned at array onset. **(C, G, K, O, S)** Mean population response aligned at fixation onset. Firing rates were normalized to the maximum response. Shaded areas denote ±SEM across units. The foveal units responded to the foveal cue stimulus but not to the peripheral stimuli in the search array before the monkeys fixated on them. In contrast, the peripheral units responded to the peripheral stimuli in the search array but not to the cue stimulus at the center of the screen. **(D, H, L, P, T)** Distribution of the category selectivity index for each group of units.

**Fig. S2.**
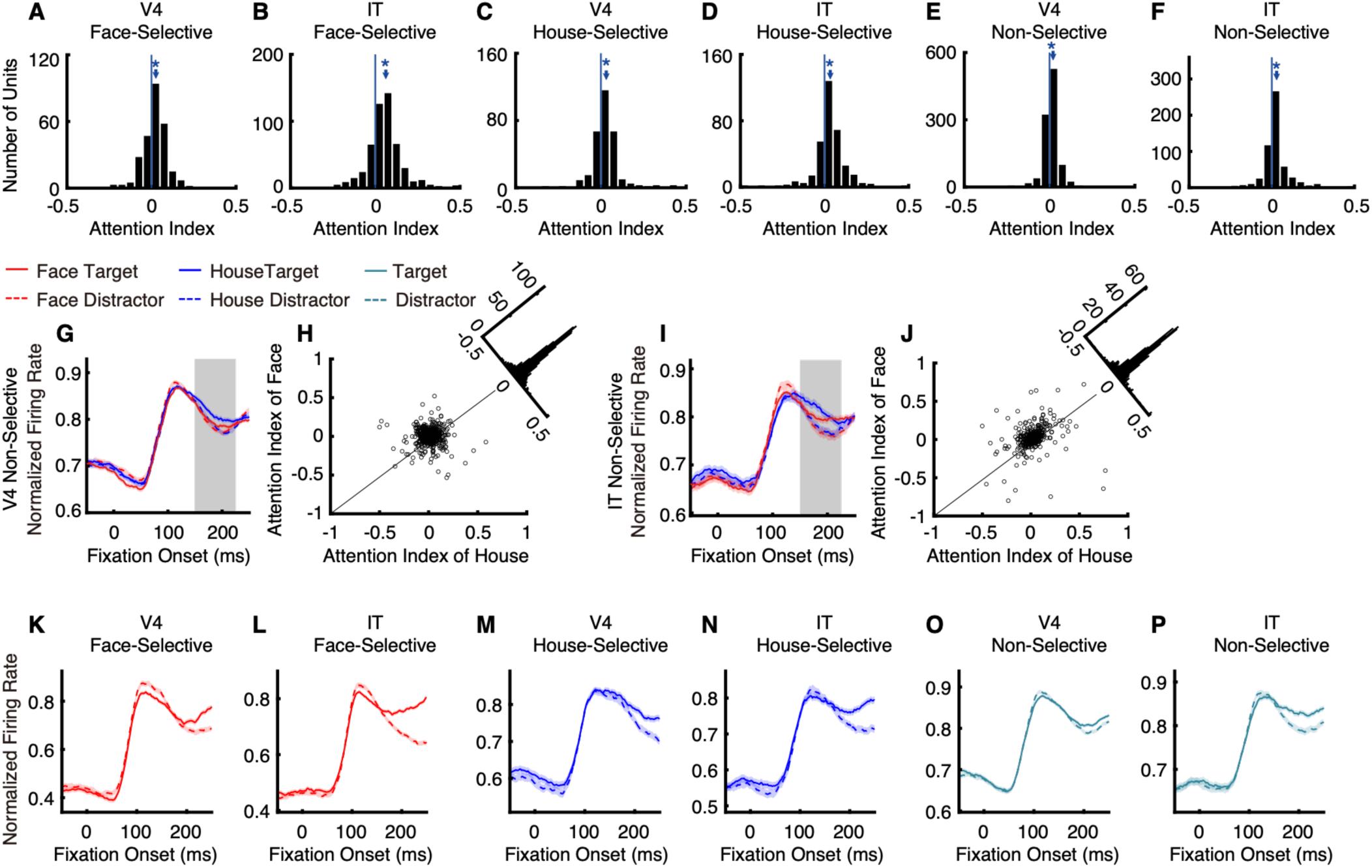
Attentional modulation in foveal units. **(A–F)** Distribution of feature attention index. **(A)** V4 face-selective units. **(B)** IT face-selective units. **(C)** V4 house-selective units. **(D)** IT house-selective units. **(E)** V4 non-selective units. **(F)** IT non-selective units. For category-selective units, we used stimuli from their preferred category. For non-category-selective units, we used both face and house stimuli. The vertical blue line indicates zero, and the blue arrow indicates the mean index. Asterisks indicate a significant difference from zero (Wilcoxon signed-rank test, *P* < 0.05). **(G, I)** Normalized population responses of foveal non-selective units during fixations on face target, face distractor, house target, and house distractor. Error shade denotes ±SEM across units. The gray shaded rectangle indicates the time window (150–225 ms after fixation onset) used to calculate the feature attention index. **(H, J)** Comparison of feature attention index for house (*x*-axis) and face (*y*-axis) stimuli in the foveal non-selective units. Each circle represents a unit. The insets at the top right show the distributions of the projections of all units onto the diagonal, which were not significantly different from zero. **(G, H)** V4 non-selective units. **(I, J)** IT non-selective units. **(K–P)** Foveal feature-based attentional modulation in V4 and IT with target and distractor fixation durations equalized. Legend conventions as in Fig. 2.

**Fig. S3.**
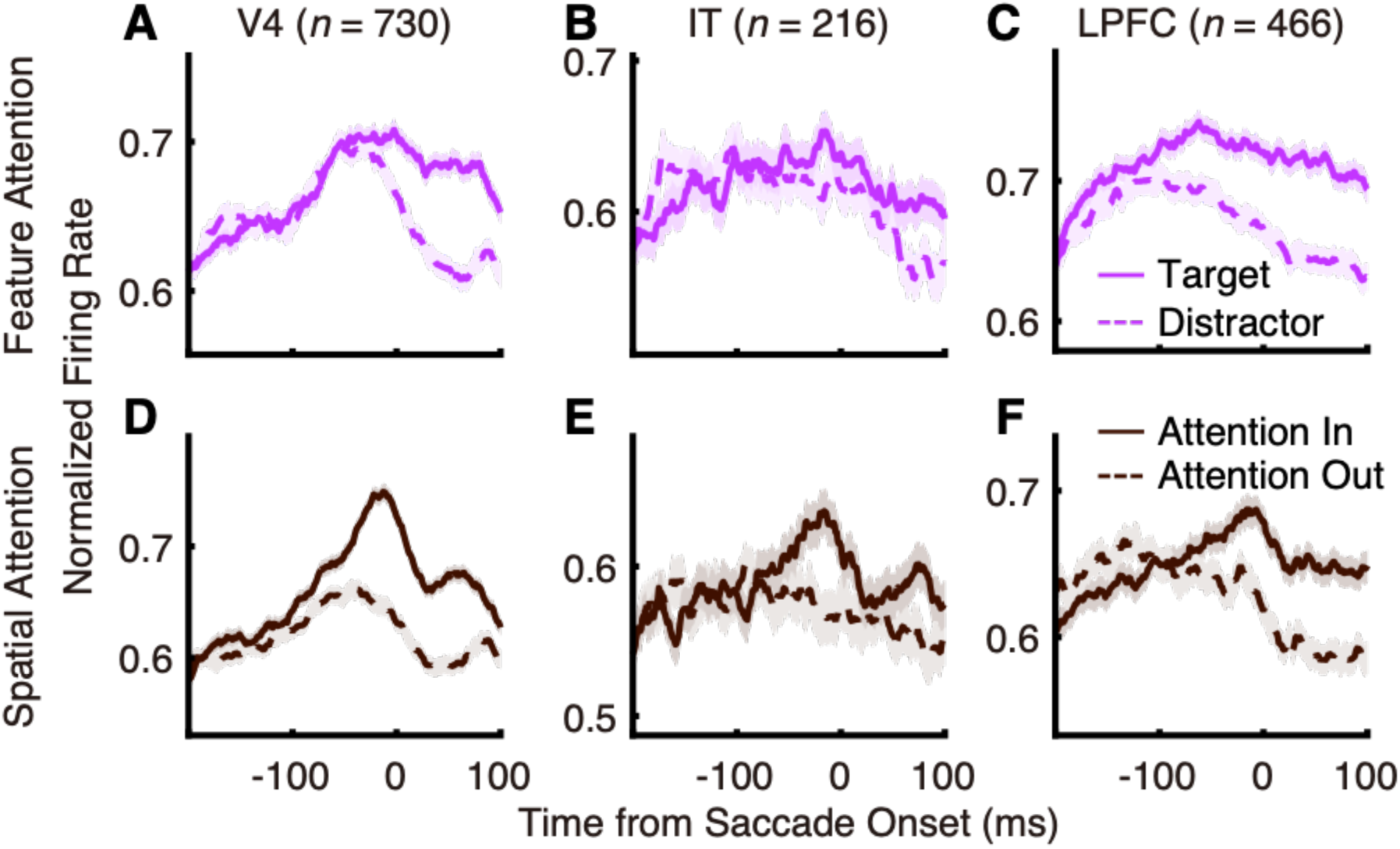
Peripheral feature and spatial attentional modulation in V4, IT, and LPFC, with responses aligned to saccade onset. Legend conventions as in Fig. 4.

**Fig. S4.**
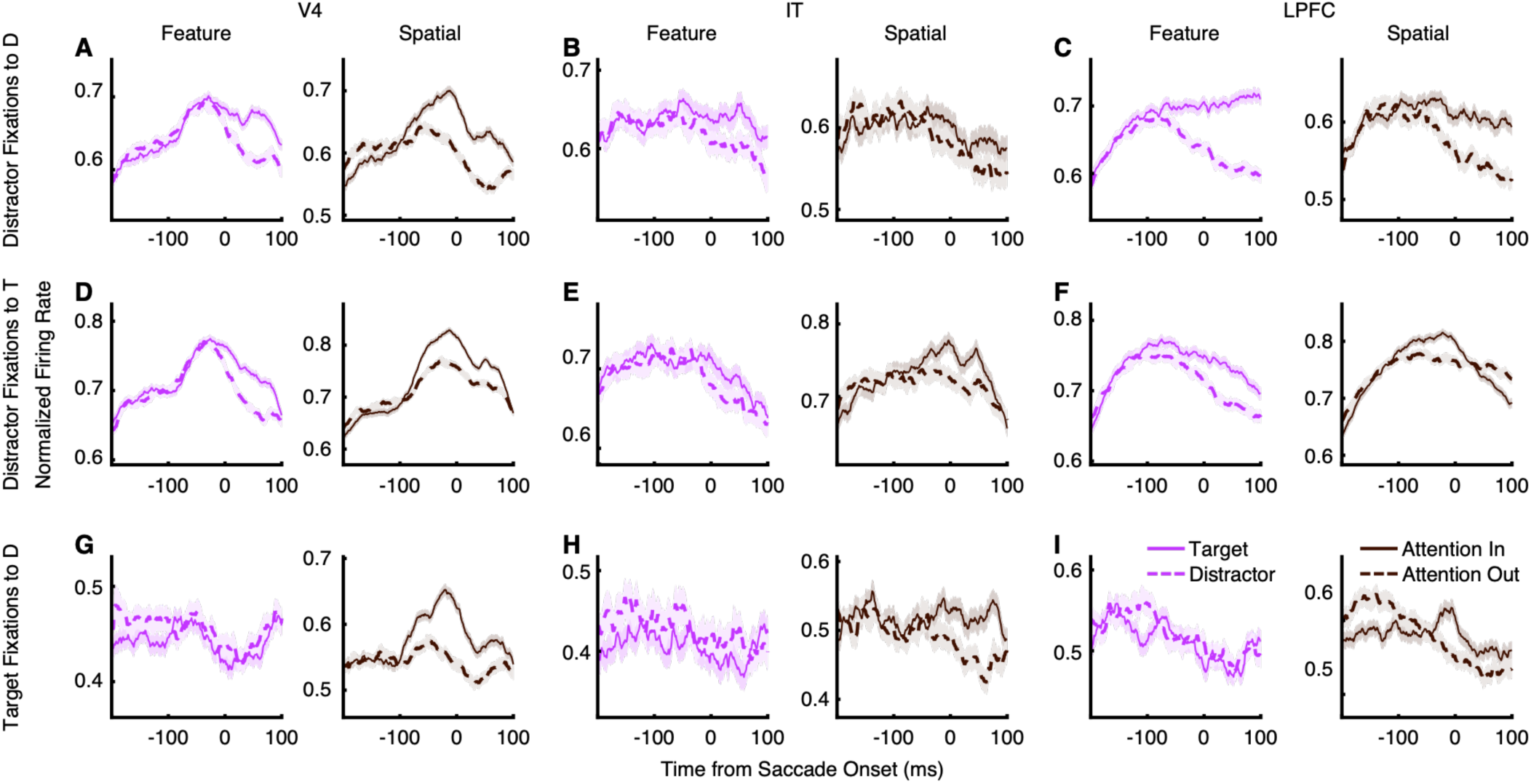
Feature and spatial attention in peripheral non-selective units, with responses aligned to saccade onset. Legend conventions as in Fig. 5.

**Fig. S5.**
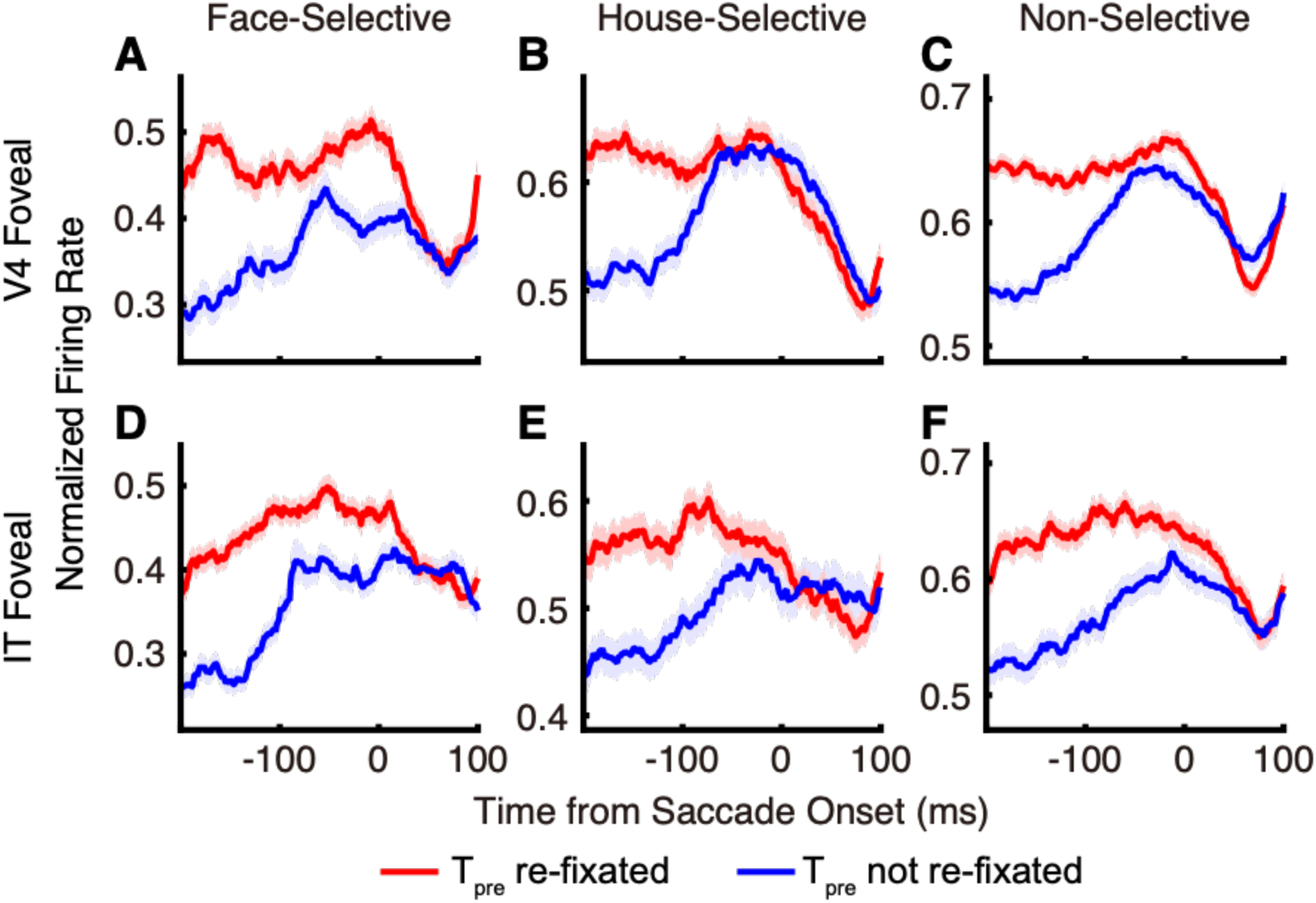
Foveal responses during saccade transitions around target fixations. Legend conventions are the same as in Fig. 6, but responses are aligned to the onset of the second saccade.

**Fig. S6.**
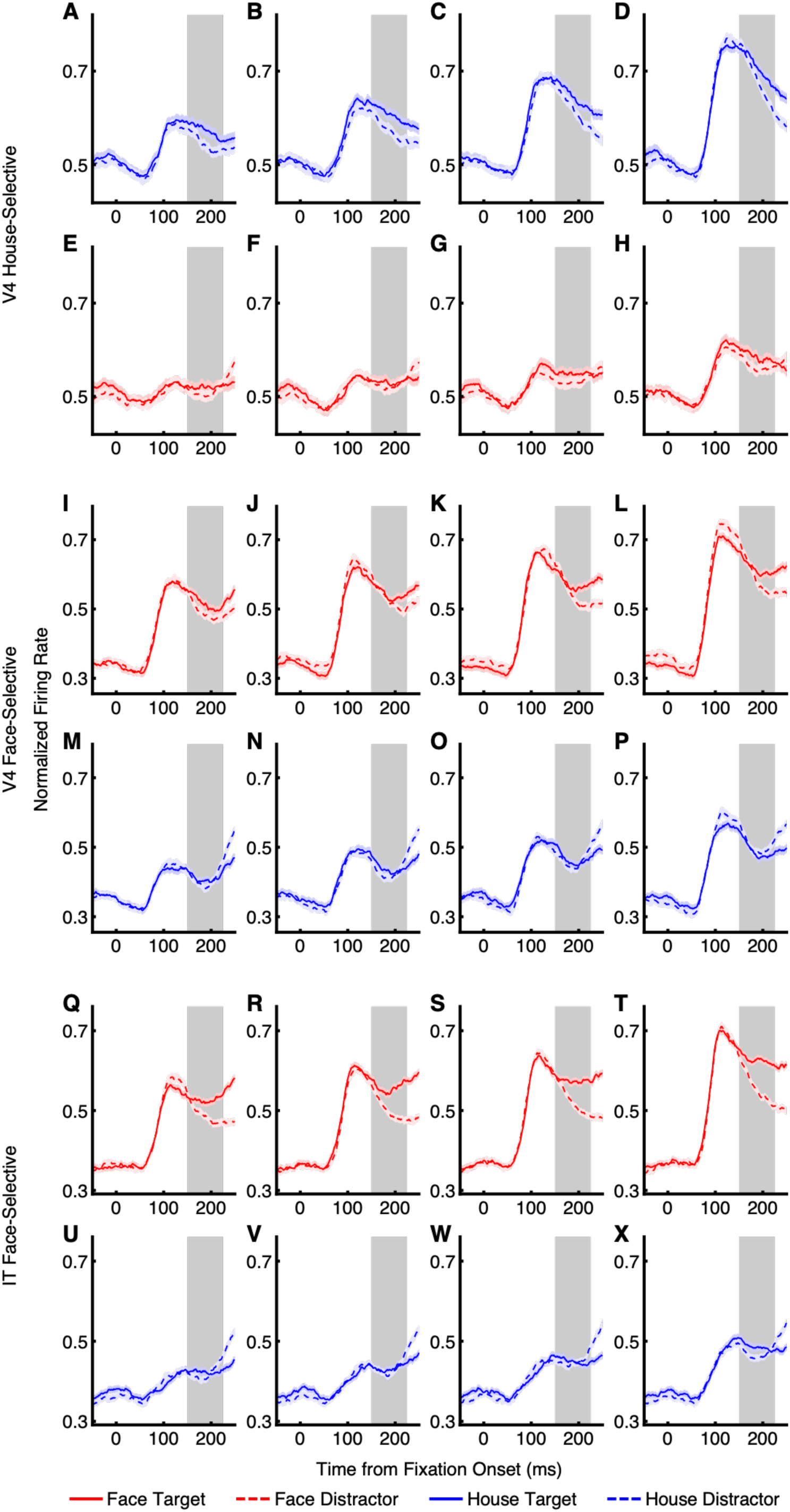
Influence of stimulus category on feature attentional modulation. Legend conventions as in Fig. 7. **(A–H)** V4 house-selective units. **(I–P)** V4 face-selective units. **(Q–X)** IT face-selective units. **(A–D, M–P, U–X)** House stimuli. **(E–H, I–L, Q–T)** Face stimuli.

